# *In Situ* Visualization of the pKM101-Encoded Type IV Secretion System Reveals a Highly Symmetric ATPase Energy Center

**DOI:** 10.1101/2021.08.19.457048

**Authors:** Pratick Khara, Liqiang Song, Peter J. Christie, Bo Hu

**Affiliations:** Department of Microbiology and Molecular Genetics, McGovern Medical School, 6431 Fannin St, Houston, Texas 77030

**Keywords:** cryoelectron tomography, DNA conjugation, type IV secretion, Pilus, protein transport

## Abstract

Bacterial conjugation systems are members of the type IV secretion system (T4SS) superfamily. T4SSs can be classified as ‘minimized’ or ‘expanded’ based on whether they are composed of a core set of signature subunits or additional system-specific components. Prototypical ‘minimized’ systems mediating *Agrobacterium tumefaciens* T-DNA transfer and pKM101 and R388 plasmid transfer are built from subunits generically named VirB1-VirB11 and VirD4. We visualized the pKM101-encoded T4SS in the native cellular context by *in situ* cryoelectron tomography (CryoET). The T4SS_pKM101_ is composed of an outer membrane core complex (OMCC) connected by a thin stalk to an inner membrane complex (IMC). The OMCC exhibits 14-fold symmetry and resembles that of the T4SS_R388_ analyzed previously by single-particle electron microscopy. The IMC is highly symmetrical and exhibits 6-fold symmetry. It is dominated by a hexameric collar in the periplasm and a cytoplasmic complex composed of a hexamer of dimers of the VirB4-like TraB ATPase. The IMC closely resembles equivalent regions of three ‘expanded’ T4SSs previously visualized by *in situ* CryoET, but differs strikingly from the IMC of the purified T4SS_R388_ whose cytoplasmic complex instead presents as two side-by-side VirB4 hexamers. Analyses of mutant machines lacking each of the three ATPases required for T4SS_pKM101_ function supplied evidence that TraB_B4_ as well as VirB11-like TraG contribute to distinct stages of machine assembly. We propose that the VirB4-like ATPases, configured as hexamers-of-dimers at the T4SS entrance, orchestrate IMC assembly and recruitment of the spatially-dynamic VirB11 and VirD4 ATPases to activate the T4SS for substrate transfer.

**SIGNIFICANCE:** Bacterial type IV secretion systems (T4SSs) play central roles in antibiotic resistance spread and virulence. By cryoelectron tomography (CryoET), we solved the structure of the plasmid pKM101-encoded T4SS in the native context of the bacterial cell envelope. The inner membrane complex (IMC) of the *in situ* T4SS differs remarkably from that of a closely-related T4SS analyzed *in vitro* by single particle electron microscopy. Our findings underscore the importance of comparative *in vitro* and *in vivo* analyses of the T4SS nanomachines, and support a unified model in which the signature VirB4 ATPases of the T4SS superfamily function as a central hexamer of dimers to regulate early-stage machine biogenesis and substrate entry passage through the T4SS. The VirB4 ATPases are therefore excellent targets for development of intervention strategies aimed at suppressing the action of T4SS nanomachines.

## INTRODUCTION

Many species of bacteria deploy type IV secretion systems (T4SSs) to deliver DNA or protein substrates to target cells (1-3). T4SSs designated as ‘minimized’ systems are assembled from a core set of signature subunits, while others termed ‘expanded’ are compositionally and structurally more complex, possibly reflecting adaptations arising over evolutionary time for specialized functions (3). In Gram-negative species, ‘minimized’ systems are assembled from ∼12 subunits named VirB1 - VirB11 and VirD4 based on the paradigmatic *Agrobacterium tumefaciens* VirB/VirD4 T4SS (3). Three subunits (VirB7, VirB9, C-terminus of VirB10) assemble as an outer membrane core complex (OMCC) that spans the distal region of the periplasm and outer membrane (OM) (4). Four integral membrane components (VirB3, VirB6, VirB8, N-terminus of VirB10) and two or three ATPases (VirB4, VirD4, +/-VirB11) together comprise the inner membrane complex (IMC) (5, 6). Some T4SSs elaborate an extracellular organelle termed the conjugative pilus from homologs of the VirB2 pilin and VirB5 pilus-tip subunit (3). ‘Expanded’ systems are composed of homologs or orthologs of most or all of the VirB/VirD4 subunits plus as many as 20 components that are system-specific (3).

To better understand the mechanism of action of T4SSs and the structural bases underlying functional diversity of this translocation superfamily, there is growing interest in solving the structures of intact machines and machine subassemblies. OMCCs are generally stable and amenable to purification, and structures are now available for OMCCs from several ‘minimized’ and ‘expanded’ systems at resolutions approaching ∼3 Å (4, 6-12). Structural analyses of inner membrane (IM) portions of T4SSs have been considerably more challenging due to problems of instability and dissociation during purification. Presently, one structure exists for a ‘minimized’ system encoded by the conjugative plasmid R388. Designated the VirB_3-10_ complex, this structure was obtained by overproduction of the VirB3 - VirB10 homologs, affinity purification of the detergent-solubilized complex, and analysis by negative-stain electron microscopy (nsEM) (6). The VirB_3-10_ complex consists of the OMCC and IMC connected by a thin, flexible stalk. The IMC is composed of a highly asymmetric IM platform connected to two side-by-side hexamers of the VirB4 ATPase extending into the cytoplasm. In an updated structure, two dimers of the VirD4 ATPase were shown to integrate between the VirB4 barrels (13).

IMCs of ‘expanded’ T4SSs have not yet been analyzed by single-particle EM. However, recent advances using *in situ* cryoelectron microscopy (CryoET) have enabled visualization of the *Legionella pneumophila* Dot/Icm, *Helicobacter pylori* Cag, and F plasmid-encoded Tra T4SSs (hereafter designated T4SS_Dot/Icm_, etc.) in the native context of the cell envelope (14-21). Remarkably, in contrast to the IMC of the VirB_3-10_ structure, the IMCs of all three ‘expanded’ systems clearly exhibit 6-fold symmetry and the VirB4 ATPases assemble as a central hexamer of dimers at the channel entrance (16-18).

Here, we solved the *in situ* structure of the pKM101-encoded T4SS, which is phylogenetically (**Fig. S1A**) and functionally closely related to the R388-encoded T4SS, to the extent that the two ‘minimized’ systems can translocate each other’s plasmids and some machine subunits are exchangeable (8). We report that the IMC of the *in situ* T4SS_pKM101_ adopts the 6-fold symmetry observed for the equivalent regions of the ‘expanded’ T4SSs. Most strikingly, the VirB4 homolog TraB (TraB_B4_) is arranged as a central hexamer of dimers, not the side-by-side hexameric barrels visualized for this ATPase in the purified VirB_3-10_ complex. Mutant machines lacking TraB_B4_ or VirB11-like TraG exhibit structural differences in the IMC compared with the wild-type machine suggestive of contributions of these ATPases to distinct stages of T4SS_pKM101_ machine assembly. Together, our findings support a model in which the VirB4 ATPases, configured as central hexamers of dimers at the bases of T4SSs, play key roles in several early-stage morphogenetic reactions required for machine biogenesis.

## RESULTS AND DISCUSSION

### *In situ* detection of the pKM101 nanomachine

To visualize T4SS_pKM101_ nanomachines, we deployed an *E. coli mreB minC* mutant carrying pKM101 to generate small (<300 nm in diameter) minicells (17). Minicells are ideal for *in situ* CryoET because of their small size and full metabolic capacity (22), including the ability to deliver plasmids such as F (17) or pKM101 **(Fig. S1B)** through encoded T4SSs to recipient cells. We used a high-throughput CryoET pipeline to visualize thousands of *E. coli* minicells (see **Fig. S2** for workflow). The pKM101 nanomachines were smaller and more difficult to detect than the F plasmid-encoded T4SS or other ‘expanded’ systems we have previously characterized (16-18), but we were able to detect pKM101-encoded structures among every 2 or 3 minicells examined (**Figs. 1Ai-iv, Movie S1**). Importantly, minicell preparations from the parental strain UU2834 alone lack these surface structures, confirming that the presence of pKM101 in the host strain is required for their elaboration.

**Fig. 1.**
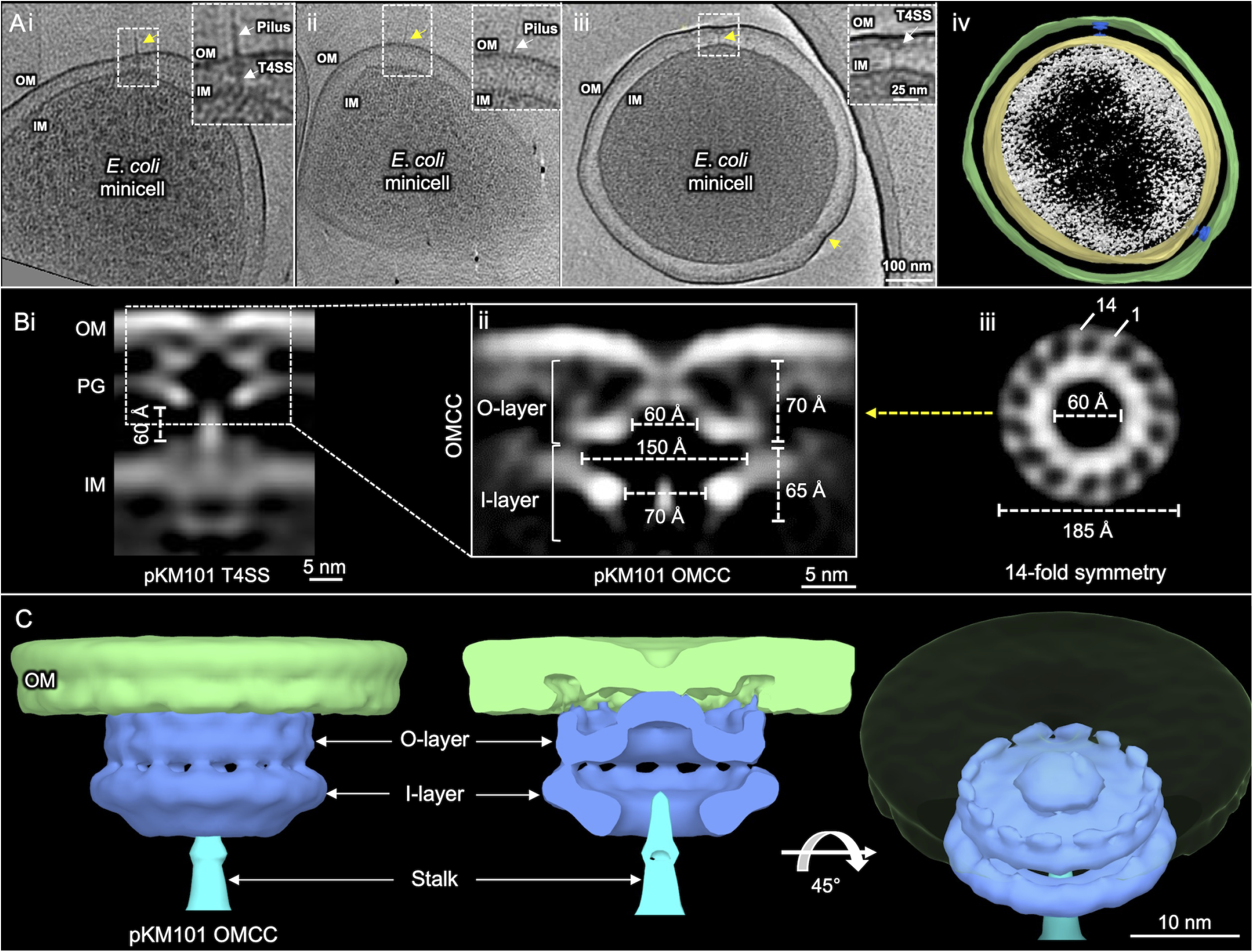
*E. coli* minicells carrying pKM101 encoded type IV secretion system (T4SS_pKM101_) and *in situ* structure of the outer membrane core complex (OMCC) of T4SS_pKM101_ revealed by CryoET and subtomogram averaging. **(Ai, ii, iii)** Tomographic slices from representative *E. coli* minicells showing T4SSs embedded between the outer membrane (OM) and inner membrane (IM). pKM101 pili were associated with a few visualized T4SSs, although pilus associated OM structures without any periplasmic densities. The T4SS and novel structures are marked with yellow arrow. The boxed regions were magnified to show T4SS with and without associated pilus and also pilus associated OM structures. **(Aiv)** A 3D surface view of the *E. coli* minicell in A iii showing T4SSs. **(Bi)** A central slice of the averaged structure of the T4SS in the cell envelope. **(Bii)** After refinement, details of the OMCC are visible. The widths and heights of O-layer and I-layer chambers are shown. **(Biii)** A cross-section view of the region in **Bii** marked by a yellow arrow shows 14-fold symmetry of the OMCC. **(C)** 3D surface renderings of the OMCC are shown in different views.

The pKM101-encoded structures consist of periplasmic cone-shaped complexes near the outer membrane (OM) without or with associated thin ‘stalk’ structures extending to the IM (**Figs. 1Ai, iii)**. OMCCs lacking stalk structures represented about half of the initially picked particles, but likely represent assembly intermediates or dead-end complexes and were not examined further (**Fig. S2**). The T4SS_pKM101_ also elaborates brittle pili that are readily detached or sloughed from cells (23). We detected some pKM101-encoded pili, bound either to the OMCC-stalk structures or to sites on the OM devoid of underlying basal densities (**Figs. 1Ai, ii, S3**). Because the pKM101 pili were rarely detected, we focused on solving the *in situ* structure of the cell-envelope-spanning nanomachine to allow for comparisons with other T4SS structures solve *in situ* or *in vitro* (4, 6, 7).

### Visualization of the *in situ* OMCC

From 287 nanomachine subtomograms extracted from 560 tomographic reconstructions, we generated *in situ* structure of the OMCC at a resolution of ∼37 Å (**Figs. 1B, S2**). Three-dimensional classifications revealed 14-fold symmetrical features of the OMCC, which were resolved further by imposing a 14-fold symmetry during refinement **(Fig. S2)**. In the refined structure, the OMCC is clearly seen attached to the OM where it causes an invagination of the outer leaflet (**Figs. 1Bi, ii)**. The upper region, designated as the O-layer (4), is 185 Å wide and 70 Å in height. In side-view, the complex forms at least two contacts with the OM, the first mediated by a central cap and the second by the periphery of the OMCC (**Fig. 1Bii**). In the middle of the central cap and extending across the OM is a region of lower density that might correspond to the OM-spanning channel. In top-down view, the periphery of the OMCC clearly consists of 14 knobs arranged in a ring of ∼185 Å in width (**Fig. 1Biii**). The knobs are connected via spokes to a central contiguous ring that conforms to the base of the cap. In 3D renderings, it is evident that the 14 peripheral knobs interact with the OM (**Fig. 1C**). Notably, besides invagination of the OM at the cap junction, the region of the OM between the central cap and peripheral contacts lacks an inner leaflet, suggesting that the OM undergoes extensive remodeling during machine biogenesis **(Figs. 1Bii, C)**.

The O-layer chamber is closed at the OM junction and widens to ∼150 Å where it joins the lower region of the OMCC known as the I-layer (4). The I-layer has a height of 65 Å and is slightly wider than the O-layer, although the outer boundary of the O-layer is blurred because of density contributed by the peptidoglycan (PG) layer **(Fig. 1Bii)**. The I-layer narrows at its base where the central chamber has a diameter of ∼70 Å. A stalk density embeds into the central cavity and projects through the periplasm to the IM (**Fig. 1Bi, ii, C**). Overall, the *in situ* OMCC structure has a total height of 135 Å (**Fig. 1Bii**).

Although the resolution of the visualized OMCC (∼37 Å) is lower than that achieved by single particle analyses (<20 Å) (4, 24), the *in situ* and *in vitro* OMCCs exhibit 14-fold symmetry and have similar cross-section dimensions of 185 Å (**Fig. S4**). They are also composed of distinct O- and I-layers that house large central chambers (4, 7). The *in vitro* structure, however, is more elongated (∼185Å in height) than the *in situ* structure (135 Å) (**Fig. S4B, C**). The O-layer and the upper portion of the I-layer are intrinsically stable due to extensive networks of interactions between the TraF_B10_ and TraO_B9_ constituents (24). By contrast, the lower portion of the I-layer, which is built from α-helical linker domains of TraF_B10_ that connect the OMCC to the IM, is highly flexible (4, 24), Therefore, we suspect that the region of the OMCC that resolves well in the *in situ* structure corresponds to the O-layer and upper portion of the I-layer, whereas the linker domains comprising the lower portion of the I-layer are either too flexible for detection *in vivo* or they fold inward to form part of the central stalk. Gratifyingly, an X-ray structure of the O-layer (7) fits well into the O-layer of the *in situ* structure (**Fig. S4D**). The OMCC of the *in vitro* VirB_3-10_ complex also generally superimposes well onto the equivalent subassembly of the *in situ* T4SS_pKM101_, although the latter structure has additional densities at the top comprising the peripheral OM contacts and laterally that might correspond to associated PG (**Fig. S4Ei - iii**), The convergence of OMCC architectures from the R388 and pKM101 systems is in line with previous findings that the OMCC from the R388 machine can be swapped for that of the pKM101 system to yield a functional chimeric system (8).

Although a flexible stalk connecting the OMCC and inner membrane complex (IMC) was previously visualized in the VirB_3-10_ complex (6), at the time it was not known if the stalk corresponded to a central channel that was structurally distorted during detergent solubilization of the nanomachine (**Fig. S4Ei, iii**). Here, our finding that a stalk density **(Fig. S4iii)** lacking a discernible channel joins the OMCC to the IMC in the *in situ* T4SS_pKM101_ confirms that the stalk is a prominent feature of ‘minimized’ systems. This distinguishes the ‘minimized’ systems from ‘expanded’ systems such as the F plasmid Tra and *L. pneumophila* Dot/Icm T4SSs (16, 17), whose *in situ* structures clearly possess central channels bridging the IMC and OMCC subassemblies (**Figs. S4Eiii, Fi-iii**). It is also interesting to note that the central stalk of the VirB_3-10_ structure spans a gap of ∼33Å between the OMCC and IMC. Here, however, we observed that the gap between the OMCCs and IMCs of different T4SS_pKM101_ machines was more variable and in the range of ∼50 - 70 Å. In situ CryoET captures structural snapshots of dynamic processes, this variability might reflect the ensemble of T4SS_pKM101_ visualized at different stages of assembly or activation.

### Visualization of the *in situ* IMC

Next, we refined the structure of the IMC using class averages of machines with detectable OMCC and IMC subassemblies (**Figs. 2A, B, S2**). Notable features of the IMC include a distinct collar surrounding the central stalk, which in top-down view presents as six knobs arranged in a ring of 155 Å. The collar was flanked by six protrusions or ‘bumps’ that were clearly distinct from the IM density (**Fig. 2B, D**). At the cytoplasmic face of the IM, the IMC was dominated by two side-by-side inverted ‘V’ structures with apices embedded in the IM and ‘arms’ projecting ∼80 Å into the cytoplasm (**Fig. 2B**). In end-on view, six V structures clearly form two concentric circles, the outer arms of the V’s configured as a knobbed ring of ∼225 Å in diameter and the inner arms joining together as a central hexameric ring with an outer diameter of ∼60 Å and a lumen of ∼45 Å. As for the periplasmic collar, the 6-fold symmetry of the concentric rings was readily visible among the class average images without symmetry imposed (**Fig. S2**). The structure was resolved further by imposing a 6-fold symmetry during refinement (**Figs. 2B-D, S2**).

**Fig. 2.**
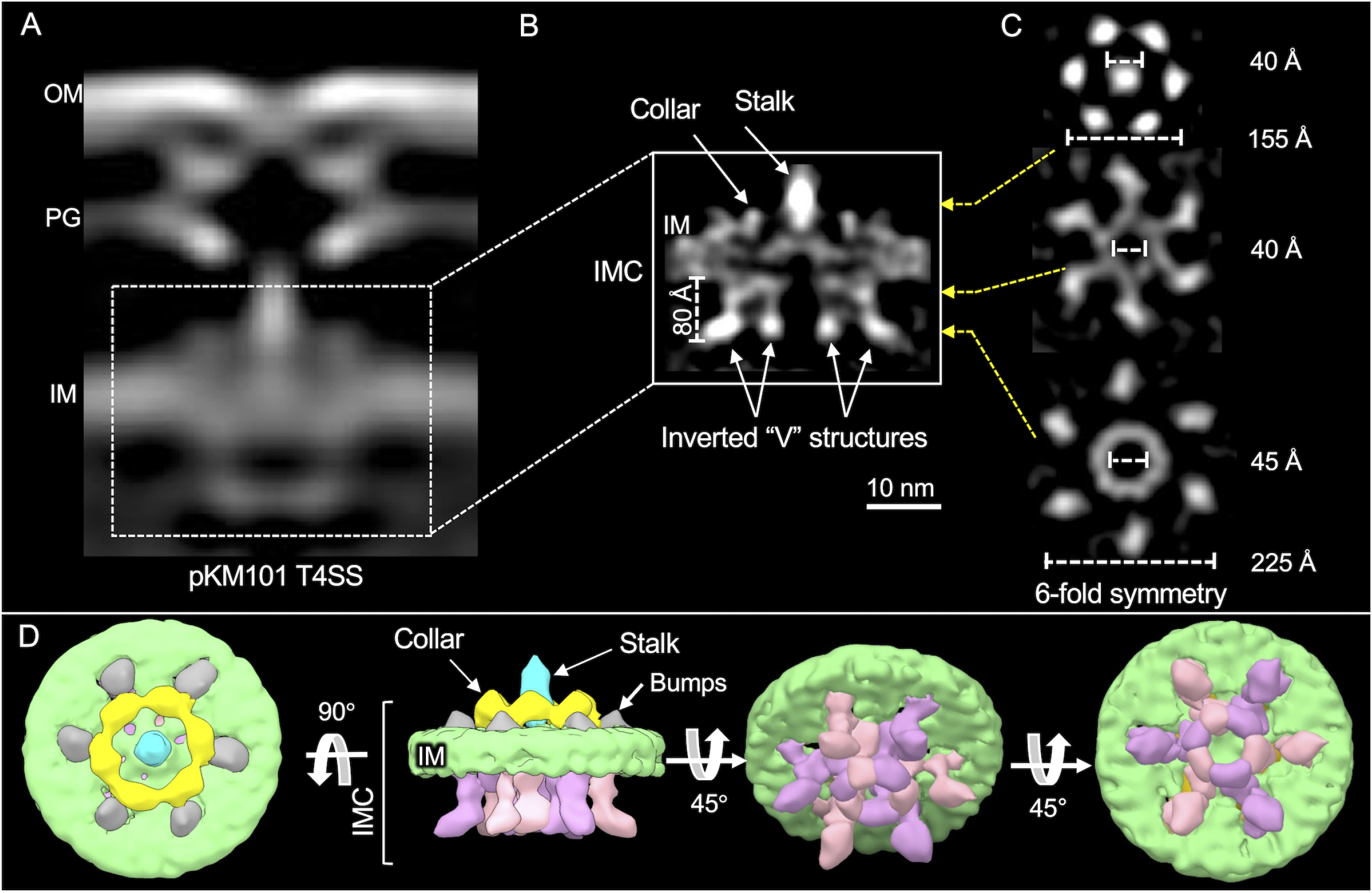
*In situ* structure of the inner membrane complex (IMC) of pKM101 encoded type IV secretion system (T4SS_pKM101_) revealed by CryoET and subtomogram averaging. **(A)** A central slice of the averaged structure of the T4SS in the cell envelope. **(B)** After refinement, details of the IMC are visible. The stalk, collar and inverted “V” structures along with the height of its arms are shown. **(C)** Cross-section views of the regions in B marked by yellow arrows show 6-fold symmetry of the IMC. The collar exists as a hexameric ring-like structure around the central stalk. **(D)** 3D surface renderings of the IMC are shown in different views.

The structure of the pKM101 IMC bears striking similarities to IMCs associated with the F plasmid-encoded Tra and *L. pneumophila* Dot/Icm systems, whose structures also were solved by *in situ* CryoET **(Fig. 3A-C)** (16, 17). Most notably, the cytoplasmic complexes of all three systems appear as side-by-side V’s in side view and as outer knobbed and inner continuous rings of similar sizes in end-on view. In studies of the F and Dot/Icm machines, structural analyses of mutant machines deleted of each of the T4SS ATPases, coupled with density tracing of a GFP moiety fused to a VirB4 homolog, established that the V structures correspond to dimers of VirB4-like ATPases (16, 17). The cytoplasmic complexes of the F plasmid and Dot/Icm systems therefore consist primarily of VirB4 subunits arranged as a central hexamer of dimers at the base of the translocation channel. The IMC of the *H. pylori* Cag T4SS is architecturally more complex than the IMCs of the F plasmid-encoded and Dot/Icm systems, yet VirB4-like Cagβ is similarly configured as a hexamer of dimers at the Cag channel entrance (18).

**Fig. 3.**
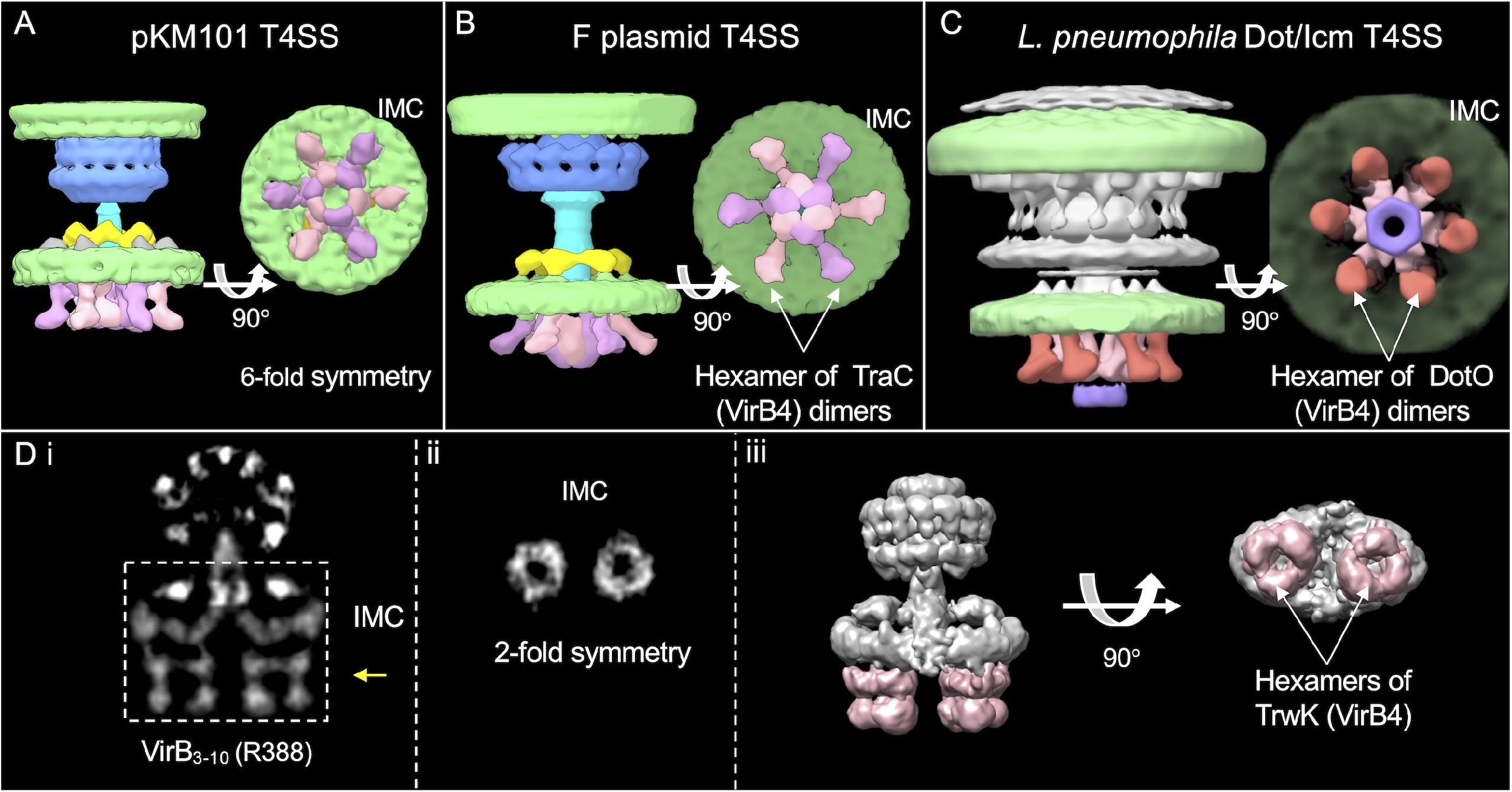
Comparison of the inner membrane complexes (IMCs) solved by CryoET and single particle analysis. **(A, B, C)** Comparison of the CryoET solved IMCs of type IV secretion systems encoded by pKM101 (T4SS_pKM101_), F plasmid (T4SS_pED208_) and *L. pneumophila* (T4SS_Dot/Icm_). 3D surface renderings showing 6-fold symmetric IMCs marked by the hexamer of dimer arrangement of VirB_4_ homologs. **(Di)** A central slice of the averaged structure of the purified VirB_3-10_ substructure encoded by plasmid R388. **(Dii)** A cross-section view of the region in **Di** marked by a yellow arrow shows 2-fold symmetry exist in IMC of purified VirB_3-10_. **(Diii)** The surface rendering of the VirB_3-10_ substructure highlighting the IMC with side-by-side two hexamers of the TrwK/VirB4 ATPase (pink-shaded). F plasmid EMD: 9344 and 9347; Dot/Icm EMD: 7611 and 7612; VirB_3-10_ EMD:2567.

The pKM101 IMC visualized here differs remarkably from that associated with the *in vitro* VirB_3-10_ structure **(Figs. 2B, C, 3A, 3Di-iii)** (6). Most notably, the pKM101 IMC is highly symmetric in its overall 6-fold symmetrical periplasmic collar and cytoplasmic complex. The VirB_3-10_ IMC is asymmetric and dominated by side-by-side barrel complexes, which consist at least partly of the VirB4 homolog TrwK as shown by gold labeling (6). A hexameric arrangement for the two barrels was inferred by earlier findings that TrwK_B4_ assembles *in vitro* as a homohexamer, and results of stoichiometric analyses showing that the VirB_3-10_ complex is composed of 12 copies of TrwK_B4_ (6, 25).

### The pKM101 cytoplasmic complex is dominated by VirB4-like TraB

To define contributions of VirB4-like TraB to the T4SS_pKM101_, we imaged Δ*traB*_*B4*_ mutant machines (100 machines from 254 tomographic reconstructions) **(Fig. S5)**. We previously reported that *traB*_*B4*_ expression in *trans* fully complements a Δ*traB*_*B4*_ mutation, confirming that the mutation is nonpolar on downstream gene expression (8). Subvolume class averages of the Δ*traB*_*B4*_ mutant machines consisted of the OMCC without associated IMC densities (**Fig. S5**). Most notably, the Δ*traB*_*B4*_ machines lacked cytoplasmic densities dominated by the concentric hexameric rings (**Fig. 4B, S5**), indicating that TraB_B4_ adopts the same hexamer-of-dimer architectures observed for VirB4 homologs associated with the F plasmid-encoded, Dot/Icm, and Cag T4SSs (16-18). This architecture is compatible with results of previous biochemical studies showing that TraB_B4_ and closely-related TrwK_B4_ from the R388 system purify as dimers or hexamers (25-28), as both of these oligomeric states are predicted from detergent extraction of a membrane ATPase with a hexamer-of-dimer configuration. The VirB4 ATPases are arranged so that their N-terminal domains (NTDs) associate tightly with the IM (28, 29), and C-terminal domains (CTDs) consisting of RecA-like α/β structural folds extend into the cytoplasm (30, 31). An atomic structure of the CTD of a VirB4 homolog fitted optimally within densities comprising the proximal halves of the V arms (**Fig. 4Eiii**), which lends further support to a conclusion that visualized hexamer of dimer densities are composed of TraB_B4_.

**Fig. 4.**
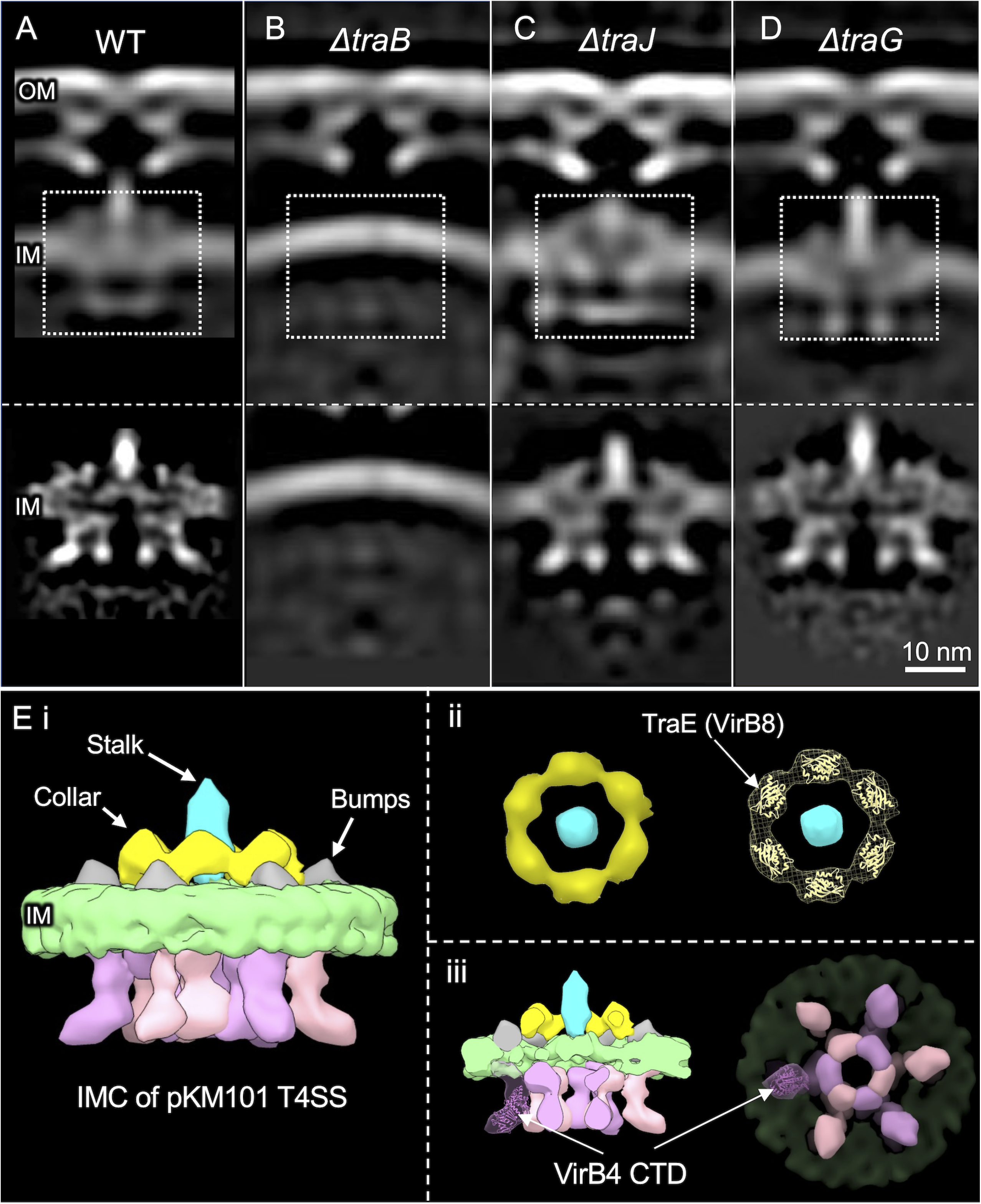
Architecture of T4SS_pKM101_ mutant machines from strains lacking one of the Tra ATPases and comparison of the IMC from T4SS_pKM101_ with those from other solved structures. **(A-D)** The top row shows central slices of the averaged structures from strains carrying native pKM101 or the Δ*traB*, Δ*traJ* and Δ*traG* mutant plasmids. The second row shows the refined IMCs of the corresponding strains. **(Ei)** A 3D surface rendering of the IMC of T4SS_pKM101_. **(Eii)** End-on view of the hexameric collar. A crystal structure of the periplasmic domain of pKM101-encoded TraE_B8_ (PDB: 5I97) fits well in the lobe-like structure of the collar. **(E iii)** A crystal structure of the C-terminal domain (CTD) of VirB4 ATPase from *Thermoanaerobacter pseudethanolicus* (PDB:4AG5) fits well in one of the arms of the TraB hexamer of dimers.

IMC densities other than the inverted V’s also were missing in Δ*traB*_*B4*_ mutant machines, including the periplasmic stalk and surrounding collar and ‘bumps’ (**Figs. 4B, S5**). VirB4 homologs associate peripherally with the IM (32) or at most possess small periplasmic domains (29), arguing against appreciable contributions of TraB_B4_ to the observed collar or stalk densities. However, several IMC subunits are strong candidates for constituting these densities. These include periplasmic linker domains of VirB10-like TraF and VirB5-like TraC, which are likely components of the stalk (6). VirB8-like TraE is a strong candidate for the central collar, as supported by a recent CryoEM structure showing that purified TraE_B8_ assembles as a homohexamer with dimensions matching those of the collar visualized *in situ* (33). Here, we also determined that an atomic structure of the periplasmic domain of TraE_B8_ (34) also fits well within each lobe of the *in situ* collar (**Fig. 4Eii**). Finally, like other VirB6 subunits (35), TraD_B6_ has a large central periplasmic domain that likely also contributes to one or more of the periplasmic densities. The absence of discernible periplasmic densities in the Δt*raB* mutant machines suggests that TraB_B4_ - IMC subunit contacts are necessary for stable assembly of the IMC. In line with this proposal, numerous studies have presented evidence that VirB4-like subunits form a network of stabilizing interactions with IMC constituents including homologs of VirB3, VirB5, VirB8, and VirB10 (8, 36-39).

### Deletions of the TraJ or TraG ATPases do not detectably alter the *in situ* T4SS_pKM101_

Nearly all T4SSs require a VirD4-like ATPase, which serves to recruit and deliver secretion substrates into the transfer channel. Designated as type IV coupling proteins (T4CPs) or substrate receptors, VirD4 subunits are members of the SpoIIIE/FtsK superfamily of motor translocases (40, 41). To determine if VirD4-like TraJ contributes to densities of the T4SS_pKM101_, we imaged Δ*traJ*_*D4*_ mutant machines (183 machines from 430 tomographic reconstructions). As observed with the WT machines, subvolume averaging yielded classes of Δ*traJ*_*D4*_ mutant machines exhibiting only the OMCC or both the OMCC and IMC densities (**Fig. S5**). A refined structure generated from the latter classes showed no distortions compared to WT machines, insofar as the OMCC, periplasmic collar and central stalk, and cytoplasmic V structures were clearly evident **(Fig. 4C)**. TraJ_D4_ thus does not contribute detectably to the *in situ* T4SS_pKM101_ structure. The F plasmid-encoded Tra and *L. pneumophila* Dot/Icm machines similarly were unaltered upon deletion of their respective VirD4 receptors (16, 17). In a recently-updated *in vitro* structure of the R388-encoded T4SS, densities thought to correspond to one or two dimers of VirD4-like TrwB were shown to be integrated between the side-by-side VirB4 barrels (13). However, the in situ architecture of the Δ*traJ*_*D4*_ mutant machines, together with evidence that VirD4-like subunits assemble as homohexamers (40, 42) and engage with T4SS channels only when activated by intracellular signals such as substrate binding and ATP hydrolysis (43-47), suggests that the *in vitro* VirB_3-10_/VirD4 complexes might represent transition-state structures.

Many T4SSs also require a third ATPase designated VirB11 for substrate transfer and pilus production (37, 48). VirB11 ATPases assemble as homohexamers that co-fractionate with the cytoplasm and IM, the latter presumably in association with the T4SS (16, 49-51). To determine if TraG_B11_ contributes to the visualized IMC_pKM101_, we imaged Δ*traG*_*B11*_ mutant machines (257 machines from 537 tomograms). We were unable to detect any density losses in the cytoplasmic complex of the Δ*traG* mutant machine compared with the WT machine (**Fig. 4**). This suggests that TraG_B11_ might associate dynamically with the T4SS_pKM101_, as shown previously for VirB11-like DotB in the *L. pneumophila* Dot/Icm system (16). In that system, detection of DotB_B11_ at the base of DotO_B4_ required deployment of a mutant form of DotB_B11_ capable of binding but not hydrolyzing ATP (16).

Interestingly, we also observed that the Δ*traG*_*B11*_ mutant machines exhibited structural aberrations compared with the WT machines that were suggestive of profound effects of TraG_B11_ docking on T4SS_pKM101_ channel architecture. Recall that OMCCs were associated with IMC densities in only ^∼^50 % of the subvolume class averages of WT machines. In striking contrast, in Δ*traG*_*B11*_ machines, ∼85 % of visualized OMCCs were associated with IMC densities. Furthermore, the IMC densities were more clearly defined for the Δ*traG*_*B11*_ mutant machines compared with WT machines, and notably the central stalks were considerably elongated (**Figs. 4, S5**). These findings suggest that TraG_B11_ plays an important role in regulating the conformational status of the central stalk and IMC. Previous work has presented evidence that VirB11 functions as a switch to regulate pilus-biogenesis vs DNA-transport modes of action in the R388-encoded system (52). Furthermore, a recent *in situ* CryoET study documented structural changes in the IM upon binding of DotB_B11_ to DotO_B4_ consistent with a role for this ATPase in opening of the IM channel in the *L. pneumophila* Dot/Icm system (21). It is enticing to propose that TraG_B11_ binding with TraB_B4_ might similarly open the IM channel of the T4SS_pKM101_ and also induce structural transitions in the central stalk of importance for substrate passage to the cell exterior. Further studies examining the structural consequences of TraG_B11_ and TraJ_D4_ docking with the T4SS_pKM101_ are clearly warranted.

## Summary

CryoET has emerged as a valuable complementary approach to single-particle CryoEM studies of bacterial secretion nanomachines (53). Although current resolutions achievable with CryoET are lower than CryoEM, structural definition of machines in their native contexts enables i) validation of architectural features observed *in vitro*, ii) assessments of machine structural variability within and between species, and iii) visualization of dynamic aspects of machine biogenesis and function (3, 53, 54). Here, we presented the first *in situ* structures of a ‘minimized’ T4SS elaborated by the model conjugative plasmid pKM101. We showed that the *in vivo* T4SS_pKM101_ consists of two large substructures, the OMCC and IMC, and that the former resembles structures of equivalent complexes solved *in vitro* (4, 7, 24). We further identified specific OM contacts, supplied evidence for OM remodeling during T4SS_pKM101_ biogenesis, and visualized a central stalk similar to that detected in the isolated VirB_3-10_ complex (6). We further gained evidence for contributions of TraGB11 to assembly or configuration of the IMC and central stalk, in agreement with recent findings for DotB_B11_ in the Dot/Icm system (21). Most importantly, we showed that TraB_B4_ assembles as a central hexamer of dimers, an oligomeric conformation similar to those of VirB4 homologs associated with the F-encoded Tra, *L. pneumophila* Dot/Icm and *H. pylori* Cag T4SSs (16-18). IMCs of T4SSs are characteristically highly unstable and difficult to purify in the presence of detergents (4, 9-12, 24), raising the possibility that the side-by-side barrel arrangement described for TrwK_B4_ in the VirB_3-10_ complex (6) might be a structural artefact of machine purification.

Together with previous biochemical and structural data (21, 36, 37, 45, 55), our findings support a model in which VirB4 ATPases play critical roles in several key steps of T4SS biogenesis, as depicted in **Fig. 5. Stage I**: The intrinsically-stable OMCC assembles without contributions by VirB4 or other ATPases. This stage I reaction is supported by our *in situ* evidence that the pKM101 OMCC assembles in the absence of associated IMC densities **(Fig. S2). Stage II:** The OMCC recruits VirB4 through previously-identified interactions between the ATPase and the cell-envelope-spanning VirB10 subunit (37), and then VirB4 recruits or stabilizes other IMC components including VirB3, VirB5, VirB6, and VirB8 to yield the IMC. This stage II reaction is supported by our findings that TraB_B4_ is required for detection of the periplasmic densities including the collar, flanking ‘bumps’, and central stalk structures **(Fig. 1). Stage III:** Upon receipt of an unknown signal, VirB4 recruits the spatially-dynamic VirB11 ATPase, which in turn induces structural changes in the stalk and IMC (**Fig. 4)** of postulated importance for the transition from a pilus-generating machine to substrate translocation channel (21, 52). **Stage IV:** Finally, upon substrate docking, the VirD4 substrate receptor binds VirB4, and the three ATPases coordinate substrate delivery through the lumen of the VirB4 hexamer and into the T4SS channel (37, 48). The proposed **stage III** and **IV** reactions are supported by our analyses of the Δ*traG*_*B11*_ and Δ*traJ*_*D4*_ mutant machines and recent findings for the Dot/Icm system (16, 21). However, it is also clear that further *in situ* studies aimed at visualizing T4SSs with stably-engaged VirB11 and VirD4 subunits, or of WT T4SSs in the act of translocating DNA or other substrates to recipient cells, will provide valuable new insights into structural transitions necessary for machine activation.

**Fig. 5.**
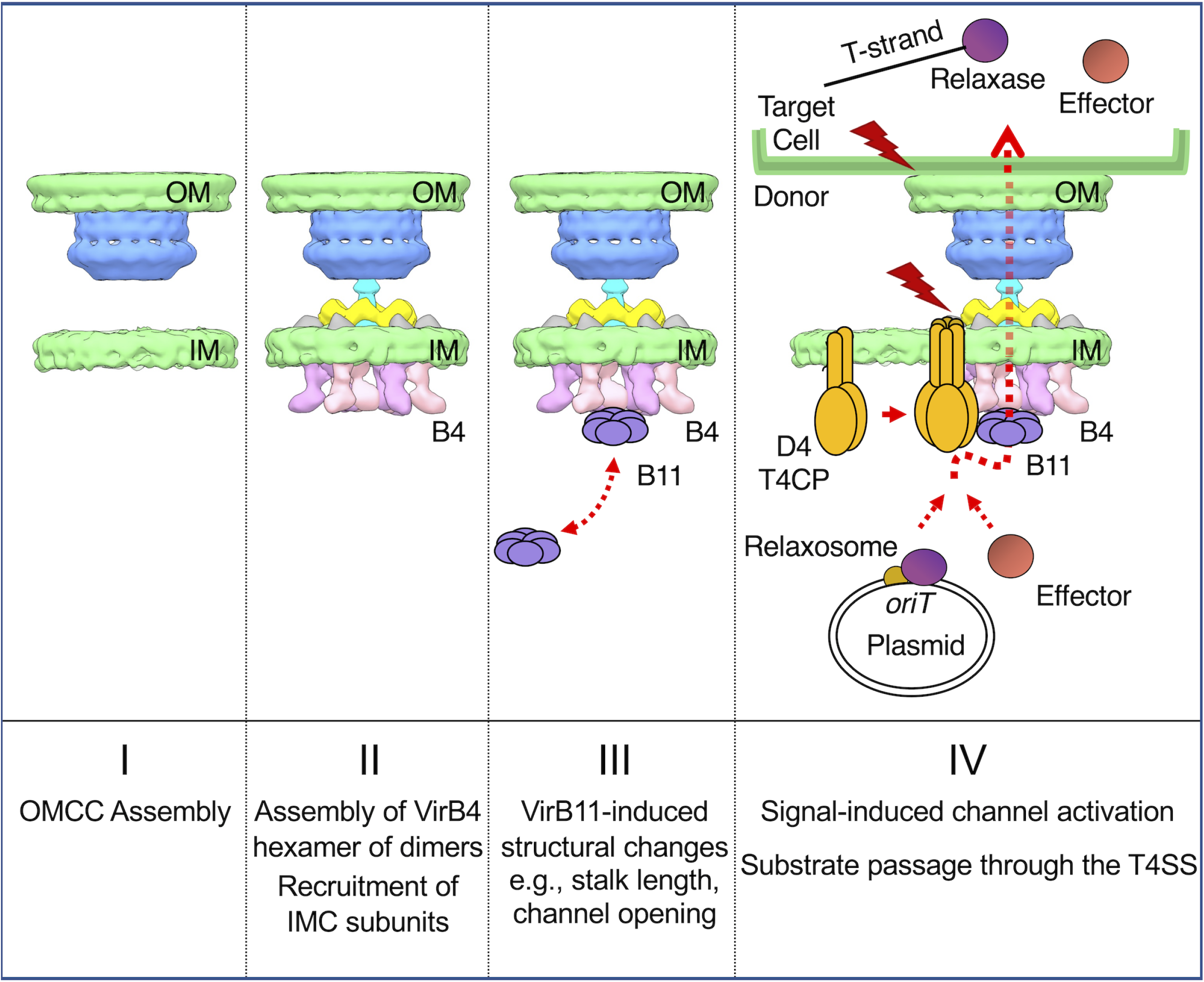
A working model depicting contributions of the VirB4 ATPases to early stages of T4SS assembly and substrate routing. **Stages: I)** The OMCC assembles as an intrinsically-stable substructure independently of contributions by the T4SS ATPases. **II)** VirB4 is recruited to the OMCC through contacts with the N-terminal cytoplasmic and IM transmembrane domains of VirB10. VirB4 assembles as a hexamer of dimers at the cytoplasmic face of the IM, where it recruits and stabilizes other IMC and stalk constituents. **III)** VirB4 serves as a docking site for the spatially-dynamic VirB11 ATPase; docked VirB11 regulates structural transitions necessary for channel activation. **IV)** The VirD4-like substrate receptor or T4CP (type IV coupling protein) oligomerizes and hydrolyzes ATP in response to binding of DNA or protein substrates; the VirD4 - substrate complex engages with and activates the T4SS channel. The ATPase energy center composed of the VirB4, VirB11, and VirD4 ATPases coordinate processing and delivery of secretion substrates through the central lumen of VirB4 and into the translocation channel (37, 48). Red lightning bolts denote signals, such as substrate binding and VirD4 ATP hydrolysis and establishment of T4SS contact with target cell, that activate the T4SS for transfer. Relaxase, the enzyme responsible for nicking the DNA strand (T-strand) destined for transfer; relaxosome, the assemblage of processing proteins at the origin-of-transfer (*oriT*) sequence that are responsible for nicking and unwinding the T-strand for transfer (see (2)).

## MATERIALS AND METHODS

### Strains and growth conditions

Bacterial strains, plasmids, and oligonucleotides used in this study are listed in Table S1. *E. coli* strains were grown at 37ºC in Luria Bertani (LB) agar or broth supplemented with appropriate antibiotics (kanamycin, 100 µg ml^-1^; spectinomycin, 100 µg ml^-1^; gentamycin, 10 µg ml^-1^). Minicells from *E. coli* strain UU2834 were used for all of the CryoET studies.

### Conjugation assays

*E. coli* MG1655 strains carrying pKM101 or mutant variants were used as donors to transfer the plasmids into UU2834 recipients. Strains containing the pKM101 mutants also harbored a complementing plasmid. Overnight cultures of donor and recipient cells grown in presence of the appropriate antibiotics at 37 ºC were diluted 1:1,000 in fresh LB media, and incubated with shaking for 1.5 h. When needed, cells were induced with arabinose (0.2 % final concentration) and incubated with shaking for another 1.5 h. Equal volumes (50 μl) of donor and recipient cell cultures were mixed and incubated for 3 h at 37ºC. Mating mixtures were serially diluted and plated onto LB agar containing antibiotics selective for transconjugants (Tc’s). Plasmid-carrying UU2834 strains were verified for the presence or absence of *tra* genes of interest by PCR. For matings to assess minicell donor capacity, minicells were spotted onto a nitrocellulose filter disc alone or with MC4100*rif*^*r*^ recipient cells, and the mating mixes were incubated at 37°C for 1 h. Discs were suspended in LB, serially diluted and plated on LB agar plates containing appropriate antibiotics selective for donors (to confirm absence of viable donor cells), recipients, or transconjugants (Tc’s). Because minicells are nonviable, the frequency of transfer is reported as Tc’s/recipient. Matings were performed two times in triplicate, and results are presented as the mean frequency of transfer with the standard error of mean (SEM) shown.

### Isolation of minicells

*E. coli* minicells were enriched essentially as described previously (56, 57). *E. coli* UU2834 harboring pKM101 or variants were grown overnight at 37°C in LB in the presence of spectinomycin, and then subcultured (1:100) in fresh LB devoid of antibiotics at 37°C to an OD_600_=0.5. Anucleate minicells were selectively enriched by centrifugation at 2,000*xg* for 10 min at room temperature to pellet rod-shaped cells. Next, the supernatant was centrifuged at 10,000*xg* for 10 min to pellet the minicells. The minicells were resuspended in fresh LB and incubated at 37°C with gentle shaking for 45 min to reinitiate cell growth. Ceftriaxone (final concentration 100 µg ml^-1^) was added to the minicell preparation to lyse growing cells, and the culture was further incubated at 37°C for 1 hr. The preparation was centrifuged at 400*xg* for 10 min to remove dead cells and debris. The supernatant was centrifuged at 10,000*xg* for 10 min to harvest the minicells. The minicells were washed twice in fresh LB, filtered through a 0.45 μm filter (Millipore) and then used for the mating assay. To minimize possible breakage of the pKM101-encoded pilus and to concentrate minicells for CryoET analyses, UU2834 strains were grown overnight on LB agar plates at 37°C. Cells were gently scraped from the plate surface with an “L” shaped rod and resuspended in phosphate-buffered saline (PBS). The cell suspension was centrifuged twice at 1,000*xg* for 3 min to remove intact cells and the supernatant was then centrifuged at 10,000*xg* for 20 min and the minicell pellet was resuspended in PBS for preparation of grids for CryoET.

### Preparation of frozen-hydrated specimens

Minicells resuspended in PBS were mixed with 10 nm diameter colloidal gold particles (Aurion BSA Gold Tracer, 10 nm) and deposited onto freshly glow-discharged, holey carbon grids (Quantifoil R2/1 200 mesh copper) for 1 min. After blotting the grids with filter paper, they were rapidly frozen in liquid ethane by using a gravity-driven plunger apparatus (58, 59).

### Cryo-ET data collection and 3D reconstructions

Frozen-hydrated specimens were imaged and data were processed using our previously established protocols (12, 28, 41). Briefly, specimens were subjected to imaging at -170°C using a Polara G2 electron microscope (FEI Company) equipped with a field emission gun and a direct detection device (Gatan K2 Summit). The microscope was operated at 300 kV at a magnification of 15K, resulting in an effective pixel size of 2.5 Å at the specimen level (17). Tomographic package SerialEM (60) was used to collect low-dose, single-axis tilt series with dose fractionation mode and a defocus at ∼6 µm and a cumulative dose of ∼60 e^-^/Å^2^ distributed over 35 stacks. Each stack contains ∼8 images. Each tilt series was collected at angles from -51° to 51° with 3° fixed increments. We used Tomoauto (58) to expedite data processing, which included drift correction of dose-fractionated data using Motioncorr (61) and assembly of corrected sums into tilt series, automatic fiducial seed model generation, alignment and contrast transfer function correction of tilt series by IMOD (62), and reconstruction of tilt series into tomograms by Tomo3D (63). Each tomographic reconstruction was 3,710 by 3,838 by 2,400 pixels and ∼130 Gb in size.

### Subtomogram averaging and correspondence analysis

Tomographic package I3 (64) was used for subtomogram analysis as described previously (65). A total of 837 T4SS_pKM101_ machines (400 × 400 × 400 voxels) were visually identified and then extracted from 1781 cryo-tomographic reconstructions. Two of the three Euler angles of each T4SS_pKM101_ machine were estimated based on the orientation of each particle in the cell envelope. To accelerate image analysis, 4 × 4 × 4 binned subtomograms (100 × 100 × 100 voxels) were used for initial alignment. The alignment proceeded iteratively, with each iteration consisting of three parts in which references and classification masks were generated, subtomograms were aligned and classified, and, finally, class averages were aligned to each other. At the initial iterations, classification mask was applied to include the whole machine and non-T4SS particles were sorted out and removed. For analysis of the IMC, a mask was applied to the IMC only, thus the T4SS particles that did not show IMC density were sorted out and the data set showing IMC was used to further refine the IMC. Classification focusing on the OMCC displayed 14-fold symmetry; therefore, 14-fold symmetry was imposed in the following processing to assist the initial alignment process. Classification focusing on the IMC showed a 6-fold symmetry feature, and in the following processing 6-fold symmetry was imposed to assist in sub-tomograms alignment. After multiple cycles of alignment and classification for 4 × 4 × 4 binned sub-tomograms, 2 × 2 × 2 binned sub-tomograms was used for refinement. Fourier shell correlation (FSC) between the two independent reconstructions was used to estimate the resolution of the averaged structures (**see Fig. S2**).

### 3D visualization

IMOD was used to visualize the maps and generate 3D surface rendering of *E. coli* minicells. UCSF Chimera (66) (http://www.rbvi.ucsf.edu/chimera) was used to visualize subtomogram averages in 3D and for molecular modeling. The video clips for the supplemental videos were made by using UCSF Chimera and further edited with iMovie.

## Data availability

Density maps and coordinate data of the T4SS_pKM101_ machines determined by cryo-electron tomography have been deposited in the Electron Microscopy Data Bank (EMDB) as EMD-24100 and EMD-24098. The authors declare that all other data supporting the findings of this study are available within the paper and its supplementary information files.

## LEGEND FOR SUPPLEMENTARY MATERIALS

**Table S1**. List of strains, plasmids, and oligonucleotides used in these studies.

**Fig. S1**. Genes and encoded functions of “minimized” type IV secretion systems (T4SSs) in Gram-negative species.

**Fig. S2**. Workflow for *in situ* CryoET.

**Fig. S3**. Detection of pKM101 pilus docked on *E. coli* outer membrane without underlying basal structures.

**Fig. S4**. Comparisons of the outer membrane core complexes (OMCCs) and stalks/channels of ‘minimized’ and ‘expanded’ T4SSs.

**Fig. S5**. Heterogeneity of the pKM101 T4SS machines revealed by class sub-volume averaging.

**Movie S1**. 3-D visualization of a tomographic reconstruction and the T4SS_pKM101_ in *E. coli* minicells.

**Movie S2**. 3-D visualization of the T4SS_pKM101_ showing architectural features of the OMCC and IMC and its comparison with purified VirB_3-10_ complex from R388.

## ACKNOWLEDGEMENTS

B.H. was supported by McGovern Medical School start-up funds, the Welch Foundation (AU-1953-20180324), NSF grant #1902392, and NIH 1R35GM138301. P.J.C. was supported by NIH 1R35GM131892 and R01GM48746. B. H. and P.J.C were supported by NIH R21AI142378. We thank members of the Christie and Hu labs for helpful discussions, and Dr. William Margolin for advice on minicell purification.

